# Green spotted puffers can detect an almost nontoxic TTX analog odor using crypt olfactory sensory neurons

**DOI:** 10.1101/2021.09.16.460554

**Authors:** Takehisa Suzuki, Ryota Nakahigashi, Masaatsu Adachi, Toshio Nishikawa, Hideki Abe

## Abstract

Tetrodotoxin (TTX) is a well-known neurotoxin that functions as a defense substance for toxic puffers. Several behavioral studies reported that TTX attracts toxic puffers belonging to the genus *Takifugu*. Although our electrophysiological and behavioral studies showed that a TTX analog, 5,6,11-trideoxyTTX, acts as an olfactory chemoattractant for grass puffers (*T. alboplumbeus*), it is unclear whether toxic puffers are commonly attracted to 5,6,11-trideoxyTTX, and which types of olfactory sensory neurons (OSNs) detect 5,6,11-trideoxyTTX.

Here we investigated whether the green spotted puffer (*Dichotomyctere nigroviridis*), a phylogenetically different species from the grass puffer, is attracted to 5,6,11-trideoxyTTX. Administration of 5,6,11-trideoxyTTX attracted green spotted puffers, but TTX or vehicle did not. Furthermore, immunohistochemistry of the olfactory epithelium exposed to 5,6,11-trideoxyTTX with an antibody against phosphorylated ribosomal protein S6 (pS6), a neuronal activity marker, labeled oval cells with apical invagination. These oval cells were also labeled by the antibody against S100, a specific marker of crypt OSNs. Thus, our results suggest that 5,6,11-trideoxyTTX acts as an olfactory chemoattractant that is detected by crypt-type OSNs in the olfactory epithelium of green spotted puffers. Toxic puffers may use 5,6,11-trideoxyTTX as an olfactory chemoattractant involved in reproduction and parental care or as an olfactory cue of TTX-bearing organisms for effective toxification.

**Summary statement:** Behavioral and immunohistochemical experiments suggest that an almost nontoxic TTX analog, 5,6,11-trideoxyTTX, acts as an olfactory chemoattractant for green spotted puffers, and crypt-type olfactory sensory neurons detect it.

## Introduction

Tetrodotoxin (TTX) is a well-known neurotoxin that functions as a defense substance for toxic puffers. TTX is believed to be accumulated in toxic puffer’s bodies through the food chain, and is originally produced by bacteria such as *Pseudoalteromonas*, *Pseudomonas*, and *Vibrio* (Magarlamov et al., 2017; Noguchi et al., 2006). TTX has also been detected in other animals, such as flatworms, ribbon worms, crabs, and gobies (Noguchi and Arakwa, 2008). Recently, TTX-bearing animals and their DNA were detected in the intestinal contents of toxic puffers, which had been eating the egg plates of toxic flatworms containing highly concentrated TTX (Itoi et al., 2015, 2018; Okabe et al., 2019).

Several studies have reported that TTX attracts toxic puffers. For example, Matsumura (1995) reported that mature male grass puffers (*Takifugu alboplumbeus*) are attracted to TTX during the spawning season. Saito et al. (2000) also reported that juvenile tiger puffers (*T. rubripes*) were attracted to TTX, and Okita et al. (2013) reported that nose ablation eliminated the attraction of tiger puffers to TTX. Our electrophysiological and behavioral study has recently shown that an almost nontoxic TTX analog, 5,6,11-trideoxyTTX, acts as an olfactory chemoattractant for grass puffers (Noguchi et al., 2021). Research has only been conducted on toxic puffers of the genus *Takifugu*, however. Thus, it is unclear if toxic puffers are commonly attracted to 5,6,11-trideoxyTTX and which types of olfactory sensory neurons (OSNs) respond to 5,6,11-trideoxyTTX.

In this study, we demonstrated that green spotted puffers are attracted to 5,6,11-trideoxyTTX, and that crypt-type OSNs are specifically activated by 5,6,11-trideoxyTTX. These findings may suggest that attraction to 5,6,11-trideoxyTTX is a common feature of toxic puffers that may be involved in reproduction, parental care, or effective toxification.

## Materials and Methods

### Animals

Green spotted puffers (*Dichotomyctere nigroviridis*; standard body length: 1.5–3.0 cm, bodyweight: 0.5–1.5 g) were purchased from a local dealer and maintained in 60 L tanks (28°C; 20–30 fish per tank) filled with artificial brackish water (ABW; Instant Oceans® Sea-Salt, Cl^-^ 543 mM, Na^+^ 471 mM, Mg^2+^ 54.3 mM, K^+^ 10.7 mM, Ca^2+^ 10 mM Br^-^ 0.7 mM, Sr^2+^ 0.1 mM, F^-^ 0.05 mM, I^-^ 0.002 mM, CO_3_^2-^/HCO_3_^-^ 3.3 mM, pH 8.2; Instant Ocean, Blacksburg, VA) with a specific gravity of 1.010 under a 12 h light/12 h dark photoperiod. Fish were fed dried shrimp once a day. In this study, both males and females were used without distinction. All fish were maintained and used in accordance with the Nagoya University regulations on animal care and use in research.

### Odorant solutions

L-Arginine (L-Arg, Sigma-Aldrich, St. Louis, MO, USA) and TTX (for biochemistry, citrate buffered, FUJIFILM Wako Pure Chemical Corporation, Osaka, Japan) were dissolved in ultrapure water and preserved as a stock solution at −30°C. 5,6,11-trideoxyTTX was synthesized and purified according to the methods of Adachi et al. (2014). 5,6,11-trideoxyTTX was dissolved in a 0.025 N acetic acid solution (for behavioral assay) or ultrapure water (for immunohistochemistry) at 10^-3^ M and stored as a stock solution at −30°C until use.

For the behavioral experiment, stock odorant solutions were diluted with ABW containing 0.8% methylcellulose. For immunohistochemistry, all stock solutions were diluted with artificial cerebro-spinal fluid for teleosts (ACSF; NaCl 140 mM, KCl 5 mM, MgCl_2_ 1.3 mM, CaCl_2_ 2.4 mM, HEPES 10 mM, glucose 10 mM; pH 7.4).

### Behavioral experiments

An overview of the system used in the experiment is shown in Fig. 1. All behavioral experiments were performed using three green spotted puffers at a time. First, three fish were transferred to a circular test aquarium (radius = 12.5 cm, depth = 8 cm) filled with ABW (2 L) prepared from deionized water and Instant Ocean® Sea-Salt and allowed to acclimate to the experimental environment for 1 to 2 h. The test aquarium was surrounded by a cardboard partition (25 cm height) so that the experimenter was invisible to the fish.

**Figure 1:**
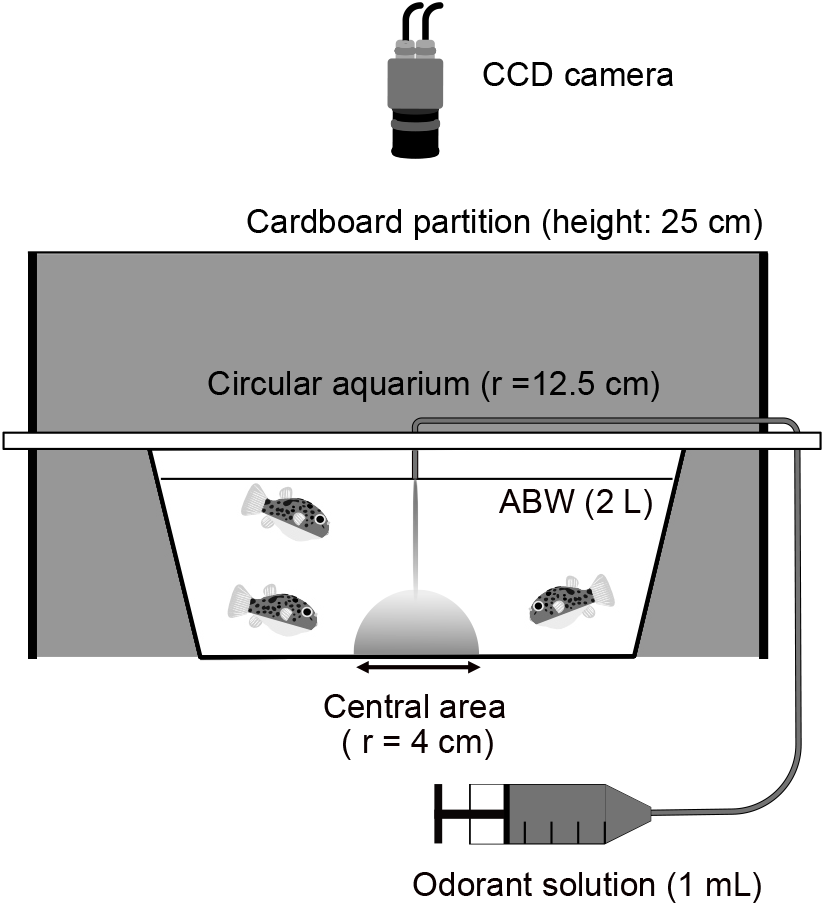
Schematic drawing of the test aquarium used in this study. Side view (left) and top view (right) of the test aquarium with odorant administration syringe. Three green spotted puffers were acclimatized in the aquarium until the odorant administration from the syringe. See the Materials and Methods section for details.

A video charge coupled device (CCD) camera (XC-ST50, Sony, Tokyo, Japan) equipped with a low-distortion C-mount lens (f = 4.4 mm; LM4NC1M, Kowa, Nagoya, Japan) was mounted vertically above the aquarium, and the camera system was adjusted so that the aquarium filled the camera frame. Time-lapse images (2 frames/s, 640 × 480 pixels) from the top of the test aquarium were acquired on a laptop computer (ThinkPad T410; Lenovo, Beijing, China) for up to 10 min using image acquisition software (Micro-Manager 1.4.22; https://micro-manager.org) and a USB video capture unit (GV-USB2, I-O Data Device, Inc., Ishikawa, Japan). Five minutes after the start of time-lapse image acquisition, 1 mL of 2 × 10^-4^ M odorant solution (vehicle [ABW or 2.5 × 10^-4^ N acetic acid in ABW], L-Arg, TTX, or 5,6,11-trideoxyTTX; diluted with ABW containing 0.8% methylcellulose) was gently administered into the center of the test aquarium using an injection needle (21G, Terumo Corporation, Tokyo, Japan) connected to a 1 mL syringe via a polyvinyl chloride tube. The estimated final odorant concentration would be 10^-7^ M if the administered solution was uniformly diffused in the test aquarium.

The numeric x-y coordinates representing the movement of the fish over time (1 coordinate per frame) were extracted from each video using UMA tracker software (http://ymnk13.github.io/UMATracker/; Yamanaka and Takeuchi, 2018). We first binarized the grayscale images using appropriate filters that the UMATracker produced. We then tracked fish position using the “Group tracker GMM” algorithm and exported the coordinates to a text file. To quantify the degree of attraction, we calculated the time spent in the central area (within a 4 cm radius of the center of the test aquarium). This criterion was based on the preliminary diffusion experiment that showed that 0.8% methylcellulose-ABW colored with methylene blue remained in that area 10 min after administration (Movie S1). We excluded the data for fish that stopped swimming for longer than 5 s during image acquisition.

### Statistics

Data are expressed as the mean ± standard error of the mean (SEM) and represented as a dot plot and a box-whisker plot (box: median and quartiles; whisker: 1.5 × interquartile range). Parametric data were analyzed using Welch’s *t*-test. Outliers were detected and excluded using Grubb’s test. Statistical analysis and graph preparations in this study were performed using R (version 4.0.4; https://www.r-project.org).

### Immunohistochemistry

Fish were anesthetized by tricaine methanesulfonate (MS–222; 0.02%, Sigma-Aldrich) and the bilateral olfactory organs were excised. The excised olfactory organs were halved using a fine blade. The olfactory epithelium (OE) was distributed on one surface of the halved olfactory organs and washed three times for 5 min in ACSF. Each OE was exposed to an odorant solution (vehicle, L-Arg, TTX, or 5,6,11-trideoxyTTX; 10^-6^ M, 1 mL, diluted with ACSF) for 5 min in a 1.5 mL test tube. After odorant stimulation, OE was fixed for 1 h in 4% paraformaldehyde at 4°C, rinsed in phosphate-buffered saline (PBS), and cryoprotected in 20% sucrose in PBS overnight at 4°C. OE was embedded in the Tissue-Tek® O.C.T. compound (Sakura Finetek Japan Co. Ltd., Tokyo, Japan). Cross cryosections (4–6 μm) of OE were mounted onto gelatin-coated slides and dried. After rehydration in PBS, the sections were blocked in 2% normal goat serum (NGS) in PBS with 0.05% Tween-20 (PBST) for 3 h at 28°C.

For single immunohistochemistry, sections were incubated successively in PBST with 1) an anti-phosphorylated ribosomal protein S6 (pS6) rabbit polyclonal antibody (Ser244, Ser247 anti-pS6, 44-923G, Thermo Fisher Scientific, MA, USA; 1:5000) that was used in mouse OSNs as validated in Sharma et al. (2017) as a neural activity marker for 2 days at 4°C; 2) an Alexa Fluor® 488-conjugated secondary antibody (goat anti-rabbit IgG (H+L), A-11008, Thermo Fisher Scientific, 1:500) for 1 day at 4°C; and 3) 4’,6-diamidino-2-phenylindole (DAPI; 1:1000 dilution) for 5 min at room temperature.

For double immunohistochemistry, sections were successively incubated with 1) a mixture of primary antibodies, anti-pS6 mouse monoclonal antibody (Ser235, Ser236 anti-pS6, 62016S, Cell Signaling Technology, Danvers, MA; 1:5000) diluted with anti-S100 rabbit serum solution (Envision FLEX-S-100, Agilent Technologies, Santa Clara, CA) that was used in zebrafish crypt OSNs as validated in Germanà et al. (2004) for 2 days at 4°C; 2) a mixture of secondary antibodies, Alexa Fluor® 555-conjugated donkey anti-mouse IgG (A-31570, Thermo Fisher Scientific; 1:500) and biotinylated goat anti-rabbit IgG (BA1000, VECTOR Laboratories, Burlingame, CA, USA; 1:500) for 1 day at 4°C; and 3) an Alexa Fluor® 488-conjugated streptavidin (S11223, Thermo Fisher Scientific; 1:500) for 2 h at 28°C. The sections were rinsed for 5 min three times in PBST after each incubation step, mounted in a mounting medium (10% Mowiol 4-88, 25% glycerol, 0.2 M Tris-HCl (pH 9.0), and 2.5%–3.0% 1,4-diazabicyclo [2,2,2] octane; diluted with ultrapure water), and cover-slipped.

### Confocal microscopy

Optical sections of the immunohistochemistry and propidium iodide–stained olfactory organs were acquired using a FV1000D IX81 confocal laser-scanning microscope (Olympus, Tokyo, Japan). All microscopic images used in this study were processed to pseudocolor stacks and were adjusted for brightness and contrast using Fiji (https://fiji.sc/). Some confocal microscopy images were analyzed using FluoRender (http://www.sci.utah.edu/software/fluorender.html; Wan et al., 2012) for three-dimensional reconstructions and visualization.

## Results

### 5,6,11-trideoxyTTX, an almost nontoxic TTX analog, attracts green spotted puffers

To investigate the behavioral response of green spotted puffers, we first created an odorant-induced behavior assay system indexed by the attraction to odorant stimuli administered to the center of a circular test aquarium. Three individuals were placed in the test aquarium and, following acclimation, L-Arg (a food odor for teleost) was administered to the center of the test aquarium (final concentration: 10^-7^ M). Before L-Arg administration, fish swam around the periphery. However, after L-Arg administration, they frequently swam into the center of the test aquarium (Fig. 2B). Vehicle administration did not change the swimming pattern of the fish (Fig. 2A). These changes in swimming patterns are shown in Movie S2. Thus, in addition to our preliminary diffusion test using the colored odorant solution, we considered the invasion to the test aquarium’s central area (within a 4 cm radius: gray circles in Fig. 2) as an indicator of attraction toward the odorant. To quantify the attractant activity, we measured the time spent on the central area every minute and compared this cumulative spending time at 5 min after L-Arg administration with that for the vehicle control (ABW). The mean cumulative spending time in the L-Arg group (23.7 ± 11.8 s, n = 6) was longer than that in the vehicle group (7.8 ± 6.5 s, n = 8) at the end of the experiment (*p* = 0.029; Fig. 3A).

**Figure 2:**
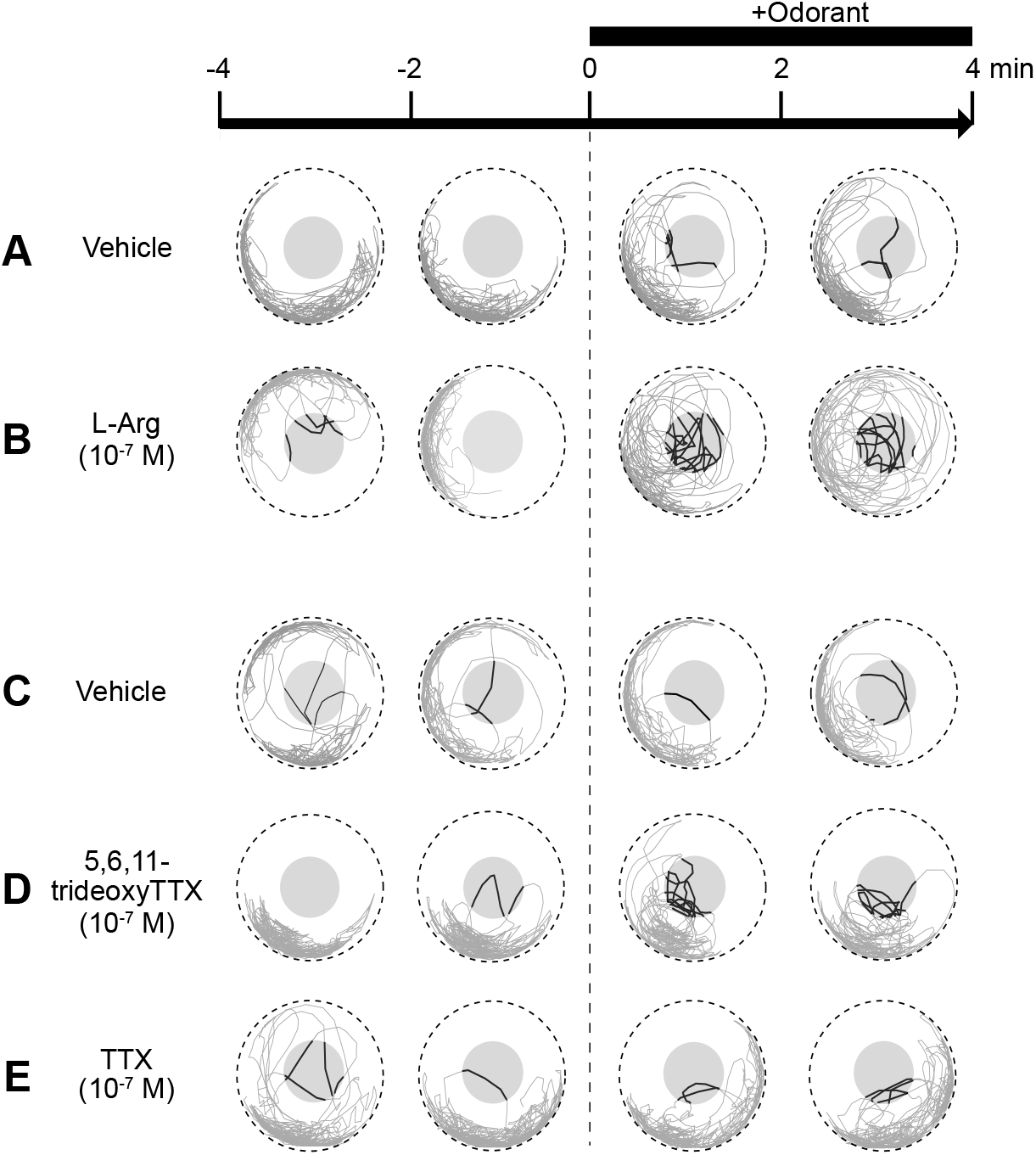
Swimming trajectories of green spotted puffers in response to odorant administration. Swimming trajectories of green spotted puffers before and after the administration of Vehicle (ABW; A), L-Arg (10^-7^ M; B), Vehicle (1.25 × 10^-7^ N acetic acids in ABW; C), 5,6,11-trideoxyTTX (10^-7^ M; D), and TTX (10^-7^ M; E). Each panel represents the swimming trajectories of three fish in 2 min under each condition; the time scale at the top show the elapsed time with the start of odorant administration as time 0. The outer circle represents the outer edge of the test aquarium, and the inner circle represents the central area within a 4 cm radius from the center of the aquarium. The swimming trajectories in the central area are shown as the bold line, and the trajectories in the other area are shown as gray lines.

**Figure 3:**
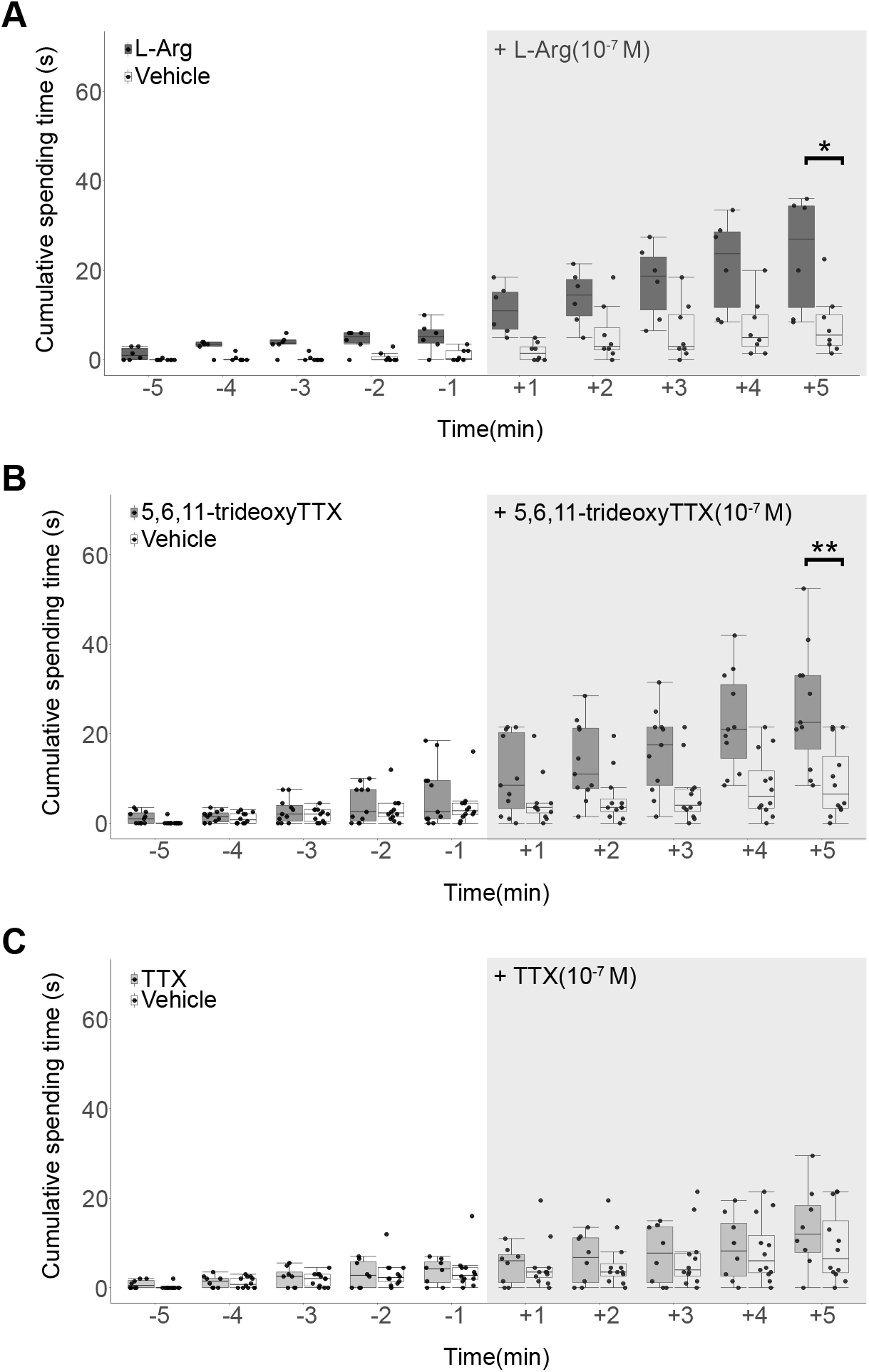
Green spotted puffers were attracted to 5,6,11-trideoxyTTX. A: Cumulative spending time of individual puffers in the central area of the test aquarium (1 min period) before and after the administration of L-Arg (10^-7^ M; gray box; n=6) or its Vehicle (ABW; open box; n=8). B: Cumulative spending time of puffers before and after the administration of 5,6,11-trideoxyTTX (10^-7^ M; gray box; n = 11) or its vehicle (1.25 × 10^-7^ N acetic acid; open box; n = 12). C: Cumulative spending time of puffers before and after the administration of TTX (10^-7^ M; gray box; n = 8) or its Vehicle (1.25 × 10^-7^ N acetic acid; open box; n = 12). The X-axis represents the elapsed time with the start of the odor administration as 0. Box plots show the median, quartiles (boxes) and 1.5 × interquartile range (whiskers). Welch’s t-test. *: *p* < 0.05, **: *p* < 0.01.

Next, we examined the changes in swimming behavior when 5,6,11-trideoxyTTX (final concentration: 10^-7^ M) or its vehicle (acetic acid in ABW; final concentration: 1.25 × 10^-7^ N) was administered. Before vehicle or 5,6,11-trideoxyTTX administration (−4 min, −2 min), green spotted puffers swam around the periphery and did not enter the central area of the test aquarium (Fig. 2C and D, respectively). However, after the 5,6,11-trideoxyTTX administrations (+2 min, +4 min), the fish actively swam in the central area (Fig. 2D, Movie S3). In contrast, the fish remained swimming peripherally after the vehicle administration and rarely invaded the central area (Fig. 2C). The cumulative spending time in the central area (1 min period) was calculated, and the time changes are plotted in Fig. 3B. Cumulative spending time increased after the 5,6,11-trideoxyTTX administration. The mean cumulative spending time of the 5,6,11-trideoxyTTX group (25.8 ± 13.0 s, n = 11) was longer than that of the vehicle group (9.4 ± 7.8 s, n = 12) at the end of the experimental period (*p* = 0.003; Fig.3B).

By contrast, when TTX was administered, the fish continued to swim around the periphery and rarely invaded the central area (Fig. 2E, Movie S4). The cumulative spending time in the central area was calculated, and the time changes are plotted in Fig. 3C. Although the mean cumulative spending time in the TTX group (13.3 ± 8.7 s, n = 8) was longer than that in the vehicle group (9.4 ± 7.8 s, n = 12), Welch’s *t*-test showed no significant difference between the two groups (*p* = 0.345). These results suggest that 5,6,11-trideoxyTTX acts as an olfactory chemoattractant in green spotted puffers as well as in grass puffers.

### 5,6,11-trideoxyTTX activated oval-shaped OSNs

The olfactory organs of teleosts are located in nasal cavities under the body surface, and their OE is located on the floor of the nasal cavity (Olivares & Schmachtenberg, 2019). Green spotted puffers have olfactory organs that protrude from the dorsal head surface with stalks and are exposed to the external environment (Fig. 4A, arrows). Each stalk is split into two fleshy flaps (Fig. 4B). The sensory epithelium (SE) is located on the inside (arrow in Fig. 4B), and the non-sensory epithelium (NSE) is located on the outside of the flap (arrowhead in Fig. 4B). Figure 4C shows a three-dimensional reconstructed confocal microscopy image of the whole-mount OE stained with propidium iodide. The inside of the flap was shallowly curved, and the SE was surrounded and separated by the NSE at the edge of the flap. OSNs are densely distributed throughout the SE. The x-z confocal section shows the cross-sectional structure of the SE (Fig. 4D). Most OSNs in the SE had spindle-shaped soma (Fig. 4D, arrow), although some large oval-shaped OSNs (minor axis diameter = c.a. 10 μm) were scattered in the SE (Fig. 4C and D, arrowheads). To identify the 5,6,11-trideoxyTTX -sensitive OSN, we immunohistochemically labeled odorant-activated OSNs using a pS6 antibody. Excised OE from the green spotted puffer was exposed to vehicle (ACSF), L-Arg, 5,6,11-trideoxyTTX, or TTX. L-Arg activated spindle-shaped OSNs (Fig. 5B, arrow). The cilia-like structure was frequently observed on the L-Arg activated OSNs (Fig. 5B inset, arrowhead). 5,6,11-trideoxyTTX activated large oval-shaped OSNs without cilia (Fig. 5C, arrow). The apical ends of 5,6,11-trideoxyTTX-sensitive OSNs were invaginated, and their cell bodies were located in the superficial layer of the OE. TTX administration weakly labeled a small number of oval-shaped OSNs (Fig. 5D, arrow). Vehicle treatment did not activate any OSNs, however (Fig. 5A). A small number of 5,6,11-trideoxyTTX-sensitive OSNs were scattered over the OE, some of which were present as cell clusters (Fig. 5E, arrows).

**Figure 4:**
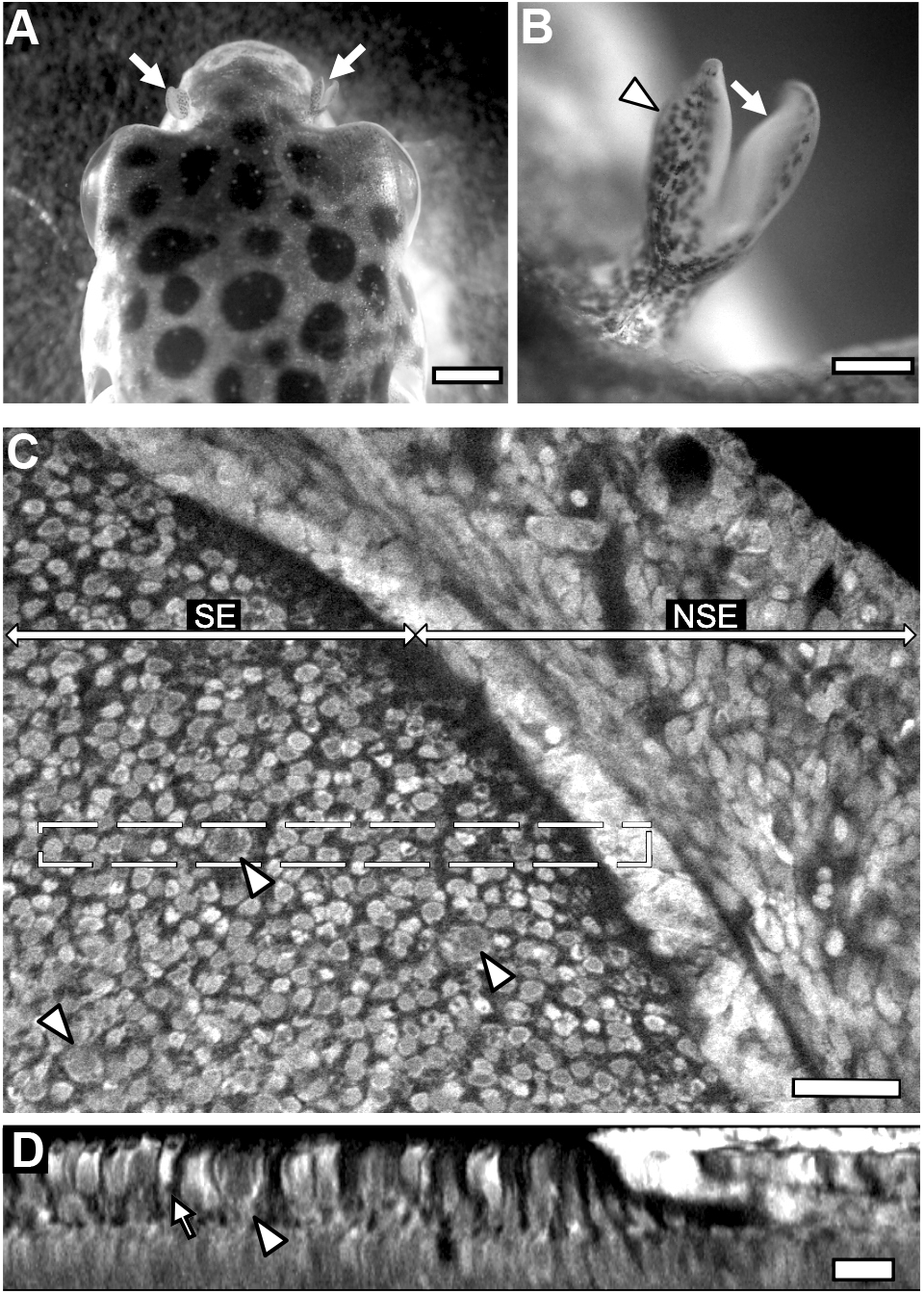
Olfactory epithelium of the green spotted puffer. A: Dorsal view of the head region of a green spotted puffer. Arrows: Olfactory organ. Scale bar: 2 mm. B: Magnified view of the olfactory organ. Arrow: Sensory epithelium, Arrowhead: Non-sensory epithelium. Scale bar: 0.5 mm. C: Reconstructed confocal microscopy image of the whole mount olfactory organ stained by propidium iodide. SE: sensory epithelium, NSE: Non-sensory epithelium. Scale bar: 20 μm. D: x-z confocal section of the dashed rectangular region in (C). Arrowhead: Large oval-shaped OSNs. Arrow: spindle-shaped OSNs. Scale bar: 10 μm.

**Figure 5:**
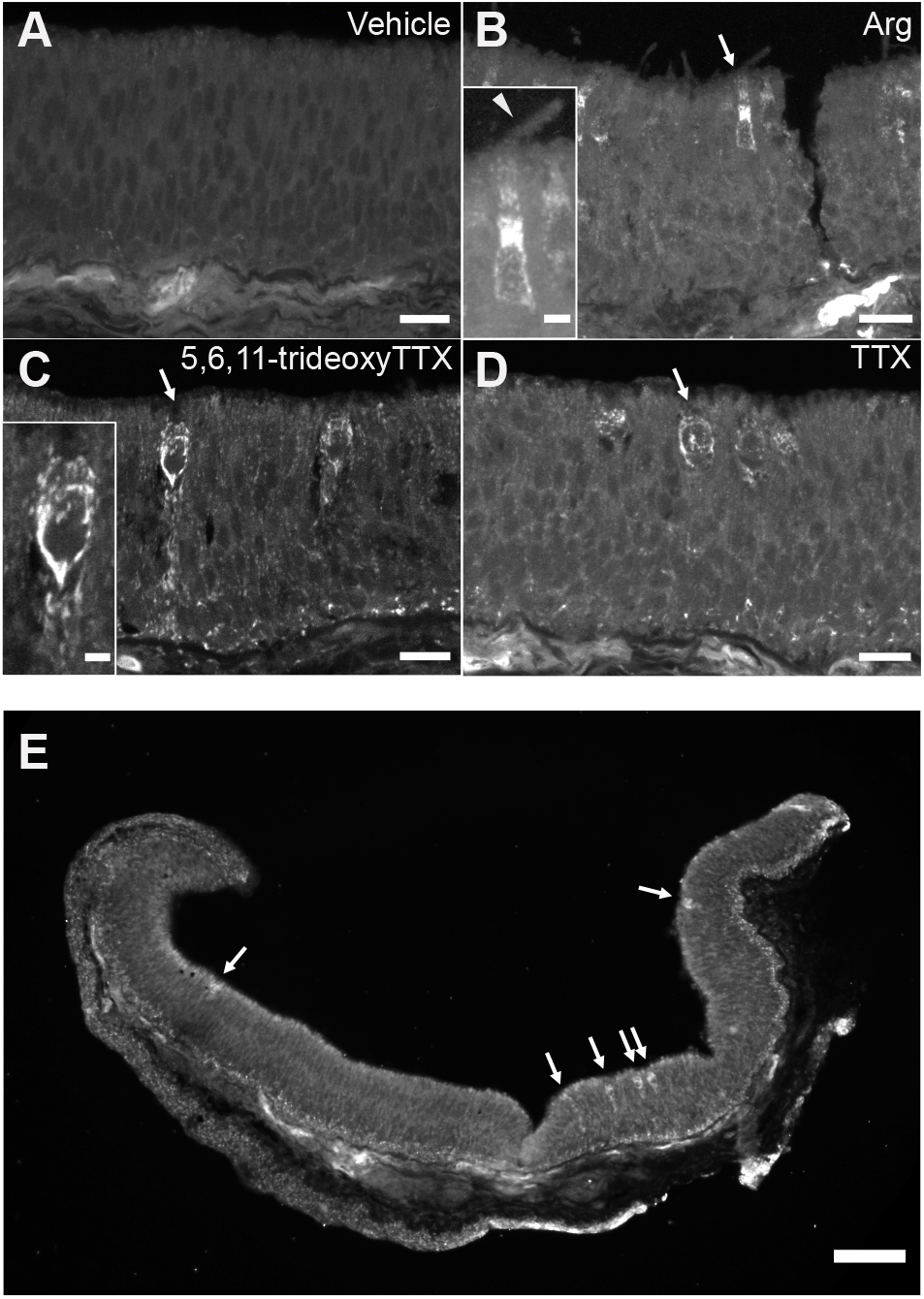
5,6,11-trideoxyTTX activates oval-shaped OSNs. A~D: pS6 immunohistochemistry of OE cross-sections exposed to Vehicle (ACSF; A), L-Arg (10^-6^ M; B), 5,6,11-trideoxyTTX (10^-6^ M; C), or TTX (10^-6^ M; D). Arrow: activated OSNs. Scale bar: 10 μm. Insets in B and C: Magnified view of pS6-immunopositive OSNs. Arrowhead: Cilia of activated OSNs. Scale bar: 2.5 μm. E: Lower magnification confocal microscopy image of pS6-immunopositive OSNs in OE cross section exposed to 5,6,11-trideoxyTTX. Arrow: 5,6,11-trideoxyTTX-activated OSNs. Scale bar: 50 μm.

### Crypt OSNs were activated by 5,6,11-trideoxyTTX

5,6,11-trideoxyTTX-sensitive OSNs were large, oval-shaped, and scattered over the OE. To identify the cellular subtype of 5,6,11-trideoxyTTX-sensitive OSNs, we performed double immunohistochemistry of 5,6,11-trideoxyTTX-treated OE with a pS6 antibody and an S100 antibody. S100 protein is a specific marker for crypt OSNs (Germanà et al., 2004). The results showed that both pS6 and S100 antibodies labeled oval-shaped OSNs with apical invagination (Fig. 6A and B). The nuclei of oval-shaped OSNs were large and located in the superficial layer of the OE (Fig. 6C, arrows). Merged confocal microscopy imaging indicated that 5,6,11-trideoxyTTX-activated pS6 immunopositive OSNs are overlapped with S100 immunopositive crypt OSNs (Fig. 6D, arrow). However, vehicle treatment did not activate any OSNs (Fig. 6E).

**Figure 6:**
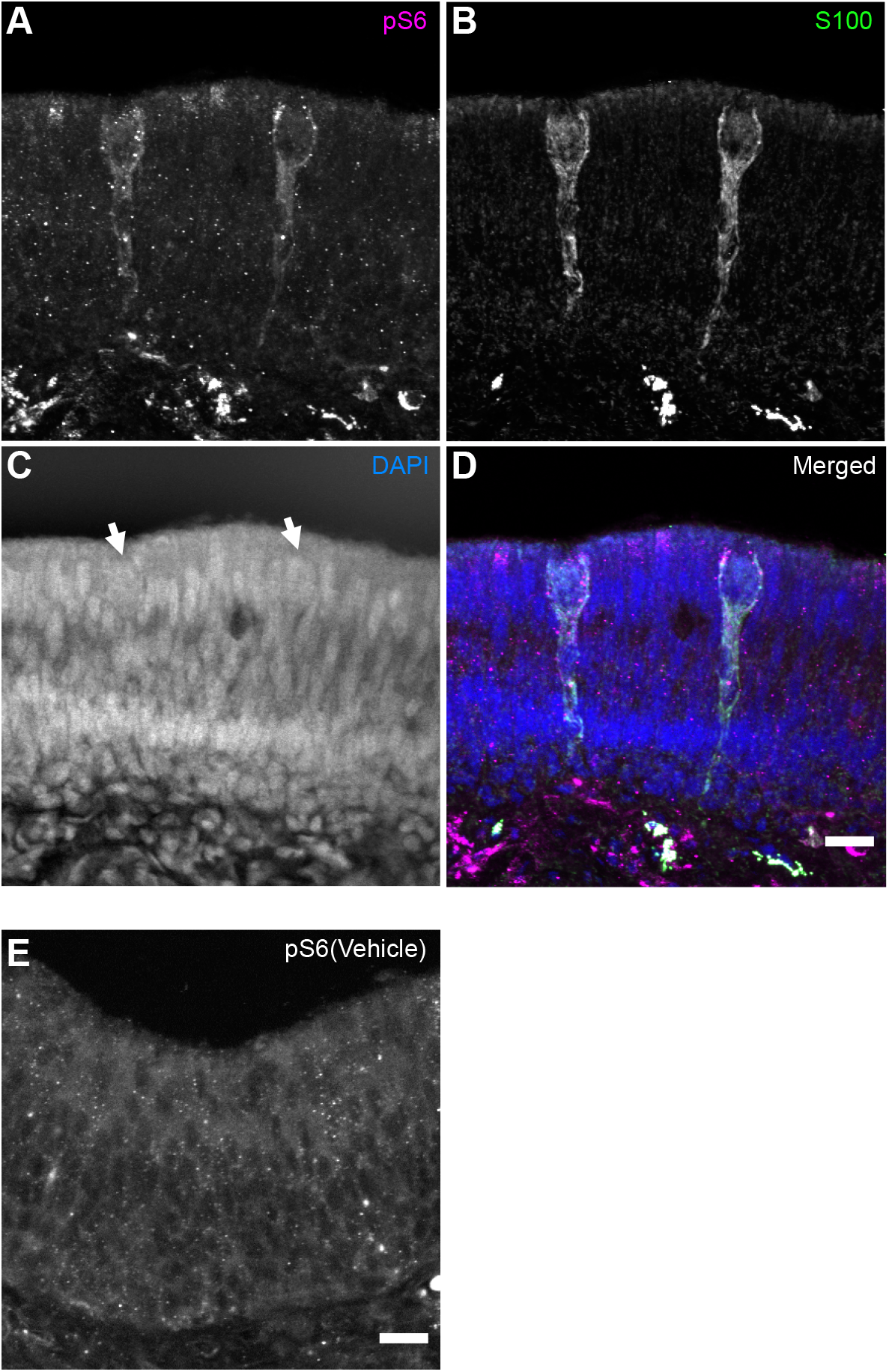
5,6,11-trideoxyTTX activates crypt OSNs. A~C: Reconstructed confocal microscopy images of 5,6,11-trideoxyTTX-administered OE displaying pS6-immunopositive OSNs (A) or S100-immunopositive OSNs (B) or stained with DAPI (C). D: A merged image of A to C (pS6: magenta, S100: green, DAPI: blue). Each immunopositive OSN had apical invagination, and their cell bodies were located in the superficial layer of the OE. Arrow: Nucleus of 5,6,11-trideoxyTTX-activated OSNs. Scale bar: 10μm. E: No pS6-immunopositive OSN was observed in Vehicle (ACSF)-treated OE. Scale bar: 10μm.

## Discussion

We demonstrated that green spotted puffers are attracted to 5,6,11-trideoxyTTX, and that crypt-type OSNs detect 5,6,11-trideoxyTTX. Previous studies reported that TTX attracts tiger puffers (*T. rubripes*, Okita et al., 2013; Saito et al., 2000) and grass puffers (*T. alboplumbeus*, Matsumura,1995; Noguchi et al., 2021). These puffers live in seawater, and the green spotted puffer lives in brackish water. Many brackish and marine Tetraodontidae species have TTX and its analogs in their body. The whole mitogenome sequence shows that Tetraodontidae can be phylogenetically divided into four clades (clade A~D; Yamanoue et al., 2011). All four clades contain both toxic and nontoxic puffers, and the genus *Takifugu* and the genus *Dichotomyctere* belong to different clades (clade C and D, respectively). Although further studies using pufferfish belonging to the other clades are needed, these phylogenic data suggest that the 5,6,11-trideoxyTTX-attracting nature appears to be a synapomorphy for marine or brackish toxic puffers belonging to these clades. The olfactory sensing ability of 5,6,11-trideoxyTTX is probably a common feature among toxic puffers.

In teleosts, there are four types of OSNs intermingled in OE: ciliated, microvillus, crypt, and kappe OSNs; the latter has not hitherto been described in teleosts other than zebrafish (Olivares and Schmachtenberg, 2019). These OSNs differ in morphology, location of cell bodies in the OE, and responsiveness to odorants (Hamdani and Døving, 2007). Ciliated OSNs are bipolar cells with a long and fine apical dendrite with a few cilia, and their cell bodies are located in the deeper layer of the OE. Microvillous OSNs are spindle-shaped bipolar cells with a short and thick apical dendrite with microvilli, and their cell bodies are located in the middle layer of the OE. Crypt OSNs are oval-shaped, have few cilia and microvilli, and their cell bodies are in the superficial layer of OE. The most striking feature of crypt OSNs is apical invaginations (Hansen and Finger, 2000). Although ciliated OSNs and microvillous OSNs are abundantly distributed, crypt OSNs are minimal and scattered in the OE.

5,6,11-trideoxyTTX-activated pS6-immunopositive OSNs were oval-shaped with apical invagination, but L-Arg-activated pS6-immunopositive OSNs were bipolar cells with cilia; thus, the shapes of each odorant-activated OSNs differ. Additionally, a small number of 5,6,11-trideoxyTTX-activated pS6-immunopositive OSNs are scattered in the OE. These morphological and locational features suggest that 5,6,11-trideoxyTTX-activated pS6-immunopositive OSNs are crypt OSNs. As such, we performed double immunohistochemistry for pS6 and S100 on the OE. Double immunohistochemistry demonstrated that 5,6,11-trideoxyTTX-activated pS6-immunopositive OSNs were S100-immunopositive and had an oval shape with apical invagination, which is a characteristic of crypt OSNs. These findings suggest that 5,6,11-trideoxyTTX-sensitive OSNs are crypt OSNs.

The functional role of crypt OSNs remains unclear. Crypt OSNs respond to the major histocompatibility complex (MHC) peptides in zebrafish (*Danio rerio*; Bienchl et al., 2016) and amino acids in mackerel (*Trachurus symmetricus*; Vielma et al., 2008). Crypt OSNs of juvenile rainbow trout respond to either amino acids, bile salts, or reproductive pheromones whereas those of mature rainbow trout respond to gonadal extracts and hormones from the opposite sex (Bazaes & Schmachtenberg, 2012). The number and density of crypt OSNs are sex-specific; moreover, the number and location of crypt OSNs in the OE varies throughout the year (Bettini et al., 2012; Hamdani et al., 2008). Studies on crucian carp showed that 1) the axons of crypt OSNs extend to the ventral part of the olfactory bulb which extends their axons to the lateral part of the medial olfactory tract (lMOT) (Hamdani & Døving, 2006), and 2) that disconnection of the lMOT reduces male reproductive behavior (Weltzien et al., 2003). Axons in the lMOT projected to the ventromedial telencephalon and the preoptic area in goldfish (Oka et al., 1982; von Bartheld et al., 1984). Moreover, electrical stimulations of the ventral telencephalon and preoptic area elicit spawning acts in male landlocked sockeye salmon, and lesions in the same areas impair courtship behavior in male goldfish (Koyama et al., 1984; Satou et al., 1984). These findings suggest that 5,6,11-trideoxyTTX-evoked signals from crypt OSNs may be involved in the reproduction of puffers.

5,6,11-trideoxyTTX has almost no toxicity and does not act as a defense substance against predators because 5,6,11-trideoxyTTX has almost no affinity to voltage-gated sodium channels (Yotsu-Yamashita et al., 1999). Toxic puffers accumulate TTX and its analogs in their bodies, but the amounts and proportions vary among species and organs. 5,6,11-trideoxyTTX is particularly accumulated in the ovaries, but the ratio of the amount of TTX and 5,6,11-trideoxyTTX in the ovaries is double in grass puffers (Jang et al., 2010). Green spotted puffers have TTX and its analogs in their liver and gonads (Chulanetra et al., 2011), and the amount of 5,6,11-trideoxyTTX in the whole body is half the amount of TTX (Jang et al., 2010). Why do toxic puffers contain this harmless toxin?

The first hypothesis is that 5,6,11-trideoxyTTX acts as an olfactory chemoattractant for conspecific puffers. Our study suggests that 5,6,11-trideoxyTTX is detected by crypt OSNs, which are thought to detect the olfactory information involved in reproduction. Matsumura (1995) reported that mature male grass puffers are attracted to the extremely low concentration of TTX (1.5 × 10^-11^ M) during the spawning season. They gather and spawn in early summer (Motohashi et al., 2010), and TTX levels in the ovaries increase during the spawning periods (Itoi et al., 2016). In addition to grass puffers, male tiger puffers chase females before spawning (Shibata et al., 2006). Although little is known about the reproductive behavior of green spotted puffers, it has been observed that their eggs are attached to hard and flat submerged surfaces in shallow water and are guarded by males until hatching; moreover, adults protect fry schools by directing them to the roots of mangroves (Pethiyagoda 1991; Rüdiger and Baensch, 1991). It has also been reported that the males of some freshwater puffers protect their eggs and use their fins to provide fresh water until they hatch. (Doi et al., 2014). These studies suggest that male green spotted puffers may use 5,6,11-trideoxyTTX to find a female for mating or to locate their eggs for parental behavior.

The second hypothesis is that toxic puffers use 5,6,11-trideoxyTTX as an olfactory cue to find TTX-bearing organisms for feeding. Recently, toxic puffer (*T. pardalis*) eggs and toxic flatworm (*Planocera multitentaculata*) DNA was detected in the intestinal contents of grass puffers, which eat the egg plates of toxic flatworms containing highly concentrated TTX (Itoi et al., 2015; Okabe et al., 2019). In another toxic puffer (*Chelonodon patoca*), DNA from toxic flatworms and gobies (*Yongeichthys criniger*) was detected in intestinal contents (Itoi et al., 2020). Furthermore, some toxic puffers secreted TTX from the glands in their skin when stimulated (Kodama et al., 1986), and predators ingested toxic puffer larva or their eggs but spat them out promptly (Itoi et al., 2014). These studies suggested that toxic puffers accumulate TTX and use it as a defense substance against predators. Because it has been reported that TTX-bearing prey organisms, such as bivalves and gastropods (Biessy et al., 2019), ribbon worms (*Cephalothrix* cf. *simula*; Vlasenko and Magarlamov, 2020), and flatworms (*Prosthiostomum trilineatum*; Suo et al., 2021) have 5,6,11-trideoxyTTX as well as TTX. Green spotted puffers feed on mollusks, crustaceans, and other invertebrates (Rainboth, 1996), and these phyla appeared to contain various TTX-bearing organisms (Noguchi and Arakawa, 2008). Thus, green spotted puffers may use 5,6,11-trideoxyTTX as an olfactory cue to selectively feed on TTX-bearing prey organisms and effectively accumulate TTX and its analogs in their body.

This study demonstrated that green spotted puffers are attracted to 5,6,11-trideoxyTTX, and that 5,6,11-trideoxyTTX activates crypt OSNs, which have been reported to sense odor information related to reproduction. These results raise the possibility that the attraction to 5,6,11-trideoxyTTX is a common characteristic among toxic puffers. Additionally, recent studies on feeding behavior suggest that 5,6,11-trideoxyTTX, as an olfactory chemoattractant, may be involved in the reproduction, parental care, or effective toxification of these toxic puffers.

## Acknowledgments

We would like to express our gratitude to Professor Naoyuki Yamamoto and Drs. Maki Goto and Hanako Hagio (Nagoya University) and Dr. Kazuya Fukuda (currently Hiroshima Univ.) for their suggestions during experimentation.

## Author contributions

Conceptualization: T.S., H.A.; Methodology: T.S., H.A., R.N., M.A., T.N.; Validation: T.S., H.A.; Formal analysis: T.S.; Investigation: T.S., R.N., H.A.; Resources: T.N., H.A.; Data curation: T.S., H.A.; Writing - original draft: T.S., H.A.; Writing - review & editing: TS, T.N., H.A.; Visualization: T.S.; Supervision: T.N., H.A.; Project administration: H.A.; Funding acquisition: H.A., T.N.

---

Movie S1: **Diffusion test of odorant solution.**

0.8%methylcellulose-ABW solution (1 mL) colored with methylene blue, which was used to dissolve the odorant solution, was administered from the center of the test aquarium filled with 2L ABW. The diffusion area of methylene blue remained within a 4 cm radius from the center of the aquarium (purple circle) even after 10 min.

Movie S2: **Attractive swimming behavior of green spotted puffers toward L-Arg.**

Vehicle or L-Arg solutions were administered from the center of the test aquarium at time 0 and remained in the aquarium. As a result, the fish frequently swam into the central area (purple circle) after L-Arg administration.

Movie S3: **Attractive swimming behavior of green spotted puffers toward 5,6,11-trideoxyTTX.**

The 5,6,11-trideoxyTTX solution was added from the center of the test aquarium at time 0 and remained in the aquarium. As a result, the fish frequently swam into the central area (purple circle).

Movie S4: **No attractive swimming behavior of green spotted puffers toward TTX.**

TTX solution was added from the center of the test aquarium at time 0 and remained in the aquarium. The fish continued to swim around peripherally and rarely invaded the central area (purple circle).

## References

Adachi, M., Sakakibara, R., Satake, Y., Isobe, M., and Nishikawa, T. (2014). Synthesis of 5,6,11-trideoxytetrodotoxin. Chem. Lett. 43, 1719–1721. doi:10.1246/cl.140684

Bazáes, A. and Schmachtenberg, O. (2012). Odorant tuning of olfactory crypt cells from juvenile and adult rainbow trout. J. Exp. Biol. 215, 1740–1748. doi:10.1242/jeb.067264

Bettini, S., Lazzari, M., and Franceschini, V. (2012). Quantitative analysis of crypt cell population during postnatal development of the olfactory organ of the guppy, *Poecilia reticulata* (Teleostei, Poecilidae), from birth to sexual maturity. J. Exp. Biol. 215, 2711–2715. doi:10.1242/jeb.069039

Biechl, D., Tietje, K., Gerlach, G., and Wullimann, M. F. (2016). Crypt cells are involved in kin recognition in larval zebrafish. Sci. Rep. 6, 24590. doi:10.1038/srep24590

Biessy, L., Boundy, M. J., Smith, K. F., Harwood, D. T., Hawes, I., and Wood, S. A. (2019) Tetrodotoxin in marine bivalves and edible gastropods: A mini-review. Chemosphere. 236, 124404. doi:10.1016/j.chemosphere.2019.124404

Chulanetra, M., Sookrung, N., Srimanote, P., Indrawattana, N., Thanongsaksrikul, J., Sakolvaree, Y., Chongsa-Nguan, M., Kurazono, H., and Chaicumpa, W. (2011). Toxic marine puffer fish in Thailand Seas and tetrodotoxin they contained. Toxins 3, 1249–1262. doi:10.3390/toxins3101249

Doi, H., Yamanoue, Y., Sonoyama, T., and Ishibashi, T. (2014). Spawning of eight Southeast Asian brackish and freshwater puffers of the genera *Tetraodon* and *Carinotetraodon* in captivity. Fish. Sci. 81, 291–299. doi:10.1007/s12562-014-0842-7

Germanà, A., G. Montalbano, R. Laurà, E. Ciriaco, M. E. del Valle, and José A. Vega. (2004). S100 protein-like immunoreactivity in the crypt olfactory neurons of the adult zebrafish. Neurosci. Lett. 371, 196–198. doi:10.1016/j.neulet.2004.08.077

Hamdani, E. H. and Døving, K. B. (2006). Specific projection of the sensory crypt cells in the olfactory system in crucian carp, *Carassius carassius*. Chem. Senses 31, 63–67. doi:10.1093/chemse/bjj006

Hamdani, E. H. and Døving, K. B. (2007). The functional organization of the fish olfactory system. Prog. Neurobiol. 82, 80–86. doi:10.1016/j.pneurobio.2007.02.007

Hamdani, E. H., Lastein, S., Gregersen, F., and Døving, K. B. (2008). Seasonal variations in olfactory sensory neurons–fish sensitivity to sex pheromones explained? Chem. Senses 33, 119–123. doi:10.1093/chemse/bjm072

Hansen, A. and Finger, T. E. (2000). Phyletic distribution of crypt-type olfactory receptor neurons in fishes. Brain Behav. Evol. 55, 100–110. doi:10.1159/000006645

Itoi, S., Ishizuka, K., Mitsuoka, R., Takimoto, N., Yokoyama, N., Detake, A., Takayanagi, C., Yoshikawa, S., and Sugita, H. (2016). Seasonal changes in the tetrodotoxin content of the pufferfish *Takifugu niphobles*. Toxicon 114, 53–58. doi:10.1016/j.toxicon.2016.02.020

Itoi, S., Kozaki, A., Komori, K., Tsunashima, T., Noguchi, S., Kawane, M., and Sugita, H. (2015). Toxic *Takifugu pardalis* eggs found in *Takifugu niphobles* gut: Implications for TTX accumulation in the pufferfish. Toxicon 108, 141–146. doi:10.1016/j.toxicon.2015.10.009

Itoi, S., Sato, T., Takei, M., Yamada, R., Ogata, R., Oyama, H., Teranishi, S., Kishiki, A., Wada, T., Noguchi, K., et al. (2020). The planocerid flatworm is a main supplier of toxin to tetrodotoxin-bearing fish juveniles. Chemosphere 249, 126217. doi:10.1016/j.chemosphere.2020.126217

Itoi, S., Ueda, H., Yamada, R., Takei, M., Sato, T., Oshikiri, S., Wajima, Y., Ogata, R., Oyama, H., Shitto, T., et al. (2018). Including planocerid flatworms in the diet effectively toxifies the pufferfish, *Takifugu niphobles*. Sci. Rep. 8, 12302. doi:10.1038/s41598-018-30696-z

Itoi, S., Yoshikawa, S., Asahina, K., Suzuki, M., Ishizuka, K., Takimoto, N., Mitsuoka, R., Yokoyama, N., Detake, A., Takayanagi, C., et al. (2014). Larval pufferfish protected by maternal tetrodotoxin. Toxicon 78, 35–40. doi:10.1016/j.toxicon.2013.11.003

Jang, J. H., Lee, J. S., and Yotsu-Yamashita, M. (2010). LC/MS analysis of tetrodotoxin and its deoxy analogs in the marine puffer fish *Fugu niphobles* from the southern coast of Korea, and in the brackish water puffer fishes *Tetraodon nigroviridis* and *Tetraodon biocellatus* from Southeast Asia. Mar. Drugs 8, 1049–1058. doi:10.3390/md8041049

Kodama, M., Sato, S., Ogata, T., Suzuki, Y., Kaneko, T., and Aida, K. (1986). Tetrodotoxin secreting glands in the skin of puffer fishes. Toxicon 24, 819–829. doi:10.1016/0041-0101(86)90107-8

Koyama, Y., Satou, M., Oka, Y., and Ueda, K. (1984). Involvement of the telencephalic hemispheres and the preoptic area in sexual behavior of the male goldfish, *Carassius auratus:* A brain-lesion study. Behav. Neural Biol. 40, 70–86. doi:10.1016/s0163-1047(84)90182-1

Magarlamov, T. Y., Melnikova, D. I., & Chernyshev, A. V. (2017). Tetrodotoxin-producing bacteria: detection, distribution and migration of the toxin in aquatic systems. Toxins 9, 166. https://doi.org/10.3390/TOXINS9050166

Matsumura, K. (1995). Tetrodotoxin as a pheromone. Nature 378, 563–564. doi:10.1017/CBO9781107415324.004

Motohashi, E., Yoshihara, T., Doi, H., and Ando, H. (2010). Aggregating behavior of the grass puffer, *Takifugu niphobles*, observed in aquarium during the spawning period. Zool. Sci. 27, 559–564. doi:10.2108/zsj.27.559

Noguchi, T. and Arakawa, O. (2008). Tetrodotoxin – distribution and accumulation in aquatic organisms, and cases of human intoxication. Mar. Drugs 6, 220–242. doi:10.3390/md20080011

Noguchi, T., Arakawa, O., and Takatani, T. (2006). TTX accumulation in pufferfish. Comp. Biochem. Physiol. Part D: Genomics Proteomics 1, 145–152. doi:10.1016/j.cbd.2005.10.006

Noguchi, Y., Suzuki, T., Matsutani, K., Nakahigashi, R., Satake, Y., Adachi, M., Nishikawa, T., and Abe, H. (2021). Almost nontoxic tetrodotoxin analog, 5,6,11-trideoxytetrodotoxin, as an olfactory chemoattractant for the grass puffers. bioRxiv doi:10.1101/2021.09.12.459881

Oka, Y., Ichikawa, M., and Ueda, K. (1982). Synaptic organization of the olfactory bulb and central projection of the olfactory tract. In Chemoreception in Fishes. (ed. T. J. Hara), pp. 61–75. Amsterdam: Elsevier

Okabe, T., Oyama, H., Kashitani, M., Ishimaru, Y., Suo, R., Sugita, H., and Itoi, S. (2019). Toxic flatworm egg plates serve as a possible source of tetrodotoxin for pufferfish. Toxins 11, 402. doi:10.3390/toxins11070402

Okita, K., Yamazaki, H., Sakiyama, K., Yamane, H., Niina, S., Takatani, T., Arakawa, O., and Sakakura, Y. (2013). Puffer smells tetrodotoxin. Ichthyol. Res. 60, 386–389. doi:10.1007/s10228-013-0353-z

Olivares, J. and Schmachtenberg, O. (2019). An update on anatomy and function of the teleost olfactory system. PeerJ 7, e7808. doi:10.7717/peerj.7808

Pethiyagoda, R. (1991). Freshwater fishes of Sri Lanka. Colombo, Sri Lanka: Wildlife Heritage Trust of Sri Lanka. pp.294 and 295.

Rainboth, W. (1996). Fishes of The Cambodian Mekong. FAO species identification field guide for fishery purposes, Rome, Italy: FAO. pp.225–227.

Rüdiger, R. and Baensch, H. A. (1991). Aquarium Atlas (vol. 1). Mergus-Verlag GmbH, Osnabrück, Germany, pp.866–867

Saito, T., Kageyu, K., Goto, H., Murakami, K., and Noguchi, T. (2000). Tetrodotoxin attracts pufferfish (“torafugu”*Takifugu rubripes)*. Bull. Inst. Ocean. Res. Dev. Tokai Univ. 21, 93–96.

Satou, M., Oka, Y., Kusunoki, M., Matsushima, T., Kato, M., Fujita, I., and Ueda, K. (1984). Telencephalic and preoptic areas integrate sexual behavior in hime salmon (landlocked red Salmon, *Oncorhynchus nerka):* Results of electrical brain stimulation experiments. Physiol. Behav. 33, 441–447. doi:10.1016/0031-9384(84)90167-7

Schmachtenberg, O. (2006). Histological and electrophysiological properties of crypt cells from the olfactory epithelium of the marine teleost *Trachurus symmetricus*. J. Comp. Neurol. 121, 113–121. doi:10.1002/cne.20847

Sharma, R., Ishimaru, Y., Davison, I., Ikegami, K., Chien, M. S., You, H., Chi, Q., Kubota, M., Yohda, M., Ehlers, M., et al. (2017). Olfactory receptor accessory proteins play crucial roles in receptor function and gene choice. eLife, 6. https://doi.org/10.7554/ELIFE.21895

Shibata, R., Aono, H., and Machida, M. (2006). Spawning ecology of tiger puffer, *Takifugu rubripes (in Japanese with English abstract)*. Bull. Fish. Res. Agen. (Japan) suppl. 4, 131–135. https://www.fra.affrc.go.jp/bulletin/bull/bull-b4/18.pdf

Suo, R., Kashitani, M., Oyama, H., Adachi, M., Nakahigashi, R., Sakakibara, R., Nishikawa, T., Sugita, H., and Itoi, S. (2021). First detection of tetrodotoxins in the Cotylean flatworm *Prosthiostomum trilineatum*. Mar. Drugs 19, 40. doi:10.3390/md19010040

Vielma, A., Ardiles, A., Delgado, L., and Schmachtenberg, O. (2008). The elusive crypt olfactory receptor neuron: Evidence for its stimulation by amino acids and cAMP pathway agonists. J. Exp. Biol. 211, 2417–2422. doi:10.1242/jeb.018796

Vlasenko, A. E. and Magalamov, T. Y. (2020). Tetrodotoxin and its analogues in *Cephalothrix cf. simula* (Nemertea: Palaeonemertea) from the sea of Japan (Peter the Great Gulf): Intrabody distribution and secretions. Toxins 12, 745. doi:10.3390/toxins12120745

von Bartheld, C. S., Meyer, D. L., Fiebig, E., and Ebbesson, S. O. E. (1984). Central connections of the olfactory bulb in the goldfish, *Carassius auratus*. Cell Tissue Res. 238, 475–487. doi:10.1007/BF00219862

Wan, Y., Otsuna, H., Chien, C. B., and Hansen, C. (2012). FluoRender: An Application of 2D Image Space Methods for 3D and 4D Confocal Microscopy Data Visualization in Neurobiology Research. In 2012 IEEE Pacific Visualization Symposium, pp. 201–208. doi:10.1109/pacificvis.2012.6183592

Weltzien, F., Höglund, E., Hamdani, E. H., and Døving, K. B. (2003). Does the lateral bundle of the medial olfactory tract mediate reproductive behavior in male crucian carp? Chem. Senses 28, 293–300. doi:10.1093/chemse/28.4.293

Yamanaka, O. and Takeuchi, R. (2018). UMATracker: an intuitive image-based tracking platform, J. Exp. Biol. 221, jeb182469. doi:10.1242/jeb.182469

Yamanoue, Y., Miya, M., Doi, H., Mabuchi, K., Sakai, H., and Nishida, M. (2011). Multiple invasions into freshwater by pufferfishes (Teleostei: Tetraodontidae): A mitogenomic perspective. PLoS ONE 6, e17410. doi:10.1371/journal.pone.0017410

Yotsu-Yamashita, M., Sugimoto, A., Takai, A., and Yasumoto, T. (1999). Effects of specific modifications of several hydroxyls of tetrodotoxin on its affinity to rat brain membrane. J. Pharmacol. Exp. Ther. 289, 1688–1696.

